# hmmIBD: software to infer pairwise identity by descent between haploid genotypes

**DOI:** 10.1101/188078

**Authors:** Stephen F. Schaffner, Aimee R. Taylor, Wesley Wong, Dyann F. Wirth, Daniel E. Neafsey

## Abstract

**Summary:** We introduce hmmIBD, software to estimate pairwise identity by decent between haploid genomes, such as those of the malaria parasite, sampled from one or more populations. We verified hmmIBD using simulated data, benchmarked it against a previously published method for detecting IBD within populations, and demonstrated its utility using *Plasmodium falciparum* data from Cambodia and Ghana.

**Supplementary information:** Supplementary data include Appendices S1, S2 and S3, and are available online.

**Availability and Implemetation:** Source code written in C99/C11-compliant C and requiring no external libraries, is freely available for download at https://github.com/glipsnort/hmmIBD/releases, alongside test datasets.

## 1 Introduction

Segments of DNA that are shared between individuals and inherited without recombination from a common ancestor are said to be identical by descent (IBD). IBD is a fundamental concept in genetics, linking ancestry to variation due to recombination, which acts on a shorter time-scale than mutation (Thompson, 2013). In the field of human genetics, IBD-based analyses are used in many different applications: to map disease loci and quantitative traits, to phase and impute genotypes, and to infer demographic histories (Thompson, 2013; Browning and Browning, 2012).

Increasingly, IBD-based analyses are also being used to study haploid organisms such as the malaria parasite. Examples include studies of malaria disease transmission (Daniels *et al.*, 2015), of malaria parasites within multiple-genotype infections (Wong *et al.*, 2017), to aid surveillance of antimalarial resistance (Cerqueira *et al.*, 2017), and to detect signals of selection (Henden,L. *et al.* (2016). *bioRxiv*). However, most existing IBD detection software (recently reviewed in Ramstetter, M.D *et al.* (2017). *bioRxiv*) is designed for humans and other diploids, and is not suitable for haploid organisms. Accordingly, malaria studies have typically used one of two programs designed for haploids: isoRelate, described in (Henden,L. et al. (2016). *bioRxiv*) and based on (Henden *et al.*, 2016), and hmmIBD, which has not been fully described in the literature to date.

In this paper, we provide a full description of hmmIBD, validate it with simulated data, and benchmark its output and performance against isoRelate. We also describe a new feature of the program, which allows inference of IBD between samples from distinct populations.

## 2 hmmIBD description

hmmIBD is based on a discrete, heterogeneous, first-order HMM with two hidden states, IBD and not-IBD; mathematical details can be found in Appendix S1. It is designed to infer IBD segments shared between pairs of haploid genomes and to estimate two quantities: (1) the marginal posterior probability of the IBD state (which can be interpreted as the expected fraction of a pair of genomes that is IBD); and (2) the rate at which the genomes transition between states, parameterized by the number of outcrossed meioses since their most recent common ancestor (MRCA), which we refer to as the number of generations. These parameters are estimated iteratively using the Baum-Welch estimation-maximization algorithm; the Viterbi algorithm is then used to calculate the most probable sequence of IBD segments (Rabiner, 1989).

Model specification requires three probability measures (Rabiner, 1989). First, initial state probabilities (IBD or not at the first position on a chromosome). These are initially set to 0.5, and then updated as the expected fraction IBD is recalculated under successive fits of the model. Second, the probabilities of changing state between two genomic positions. These state transition probabilities are functions of the distances between positions (in base pairs), the recombination rate and the number of generations since the MRCA (both assumed to be uniform across the genome), and the expected fraction IBD. Third, emission probabilities, which are the probabilities of the observed allelic types given IBD or not. These are calculated as follows. If two genomes are IBD at a given genomic position, they are of the same allelic type, meaning no mutations are assumed to have occurred since the MRCA. If they are not IBD, the alleles are modeled as independent draws based on the allele frequencies in the population. The probability of the observed allele is then calculated from these probabilities by allowing for genotyping error.

Parameters include the recombination rate (the default value in the code for *P. falciparum* (Miles *et al*., 2016)) and the genotyping error rate, as well as estimates of the allele frequencies for all variable sites in the input dataset. Allele frequencies can be supplied by the user; if not supplied, the program estimates them from the data. The model accommodates positions with missing data by omitting emission probabilities at those sites. The program is implemented in C; it complies with the Cll standard and requires no additional libraries. It can accept genotype data for any variant that can be represented by an integer at a single position on the chromosome (e.g. all SNPs and most small indels).

One important assumption of the model is that all IBD regions present between two samples result from common ancestry on a similar time-scale. Clearly, this need not be the case: very recent inbreeding can be present along with much older background sharing, the kind that generates the linkage disequilibrium in the population. Because IBD deriving from recent common ancestry is of greater interest for many applications, hmmIBD provides an option of capping the number of generations to the MRCA in the model; its effect is to bias against breaking up segments of either state.

## 3 hmmIBD performance

We verified the performance of hmmIBD using data simulated under the HMM on which it is based; details and full results can be found in S2. For a typical situation, the RMS error on the IBD fraction was 0.25 percentage points and on the number of generations was 2 generations. CPU time was linear in the number of variants. We next bench-marked hmmIBD against isoRelate (Henden,L. et al. (2016). *bioRxiv*) using data created from artificially recombined field samples; details and full results can be found in S3. Both programs performed well, with sensitivities and specificities greater than 97% (Table 1). On average, hmmIBD was 25 times faster than isoRelate. (We note that isoRelate has a unique capability: by modeling the hidden state space as a set of IBD allele counts in 0, 1 or 2, it can accommodate samples containing two genotypes.)

**Table 1.** Benchmark results of isoRelate and hmmIBD

| Software | Clock time per 50 samples (sec) | Accuracy | Sensitivity | Specificity |
| --- | --- | --- | --- | --- |
| isoRelate | 1711 (365) | 0.995<br>(0.004) | 0.999<br>(0.002) | 0.991<br>(0.008) |
| hmmIBD | 67 (12) | 0.993<br>(0.005) | 0.999<br>(0.001) | 0.986<br>(0.011) |

An unusual feature of hmmIBD is that it can accommodate samples from different populations, even ones with very different allele frequencies. We believe this feature can have multiple applications, including detection of selective sweeps that spread between populations and determining the source of imported cases in elimination settings. We demonstrate its effectiveness by examining IBD in the region of the selective sweep for chloroquine resistance in *P. falciparum* around the gene *pfcrt*. Figure 1 shows amount of IBD between field isolates from Ghana and those from Cambodia. The increase in IBD around *pfcrt* is clear, reflecting the fact that the resistance haplotype (Wellems *et al*., 1991) emerged and spread into Africa from South-East Asia (Wootton *et al*., 2002; Ariey *et al*., 2006). An alternative approach using hmmIBD but treating the samples as coming from a single population as in (Henden,L. et al. (2016). *bioRxiv*), also shown, is much less effective at detecting the cross-population IBD.

**Fig. 1.**
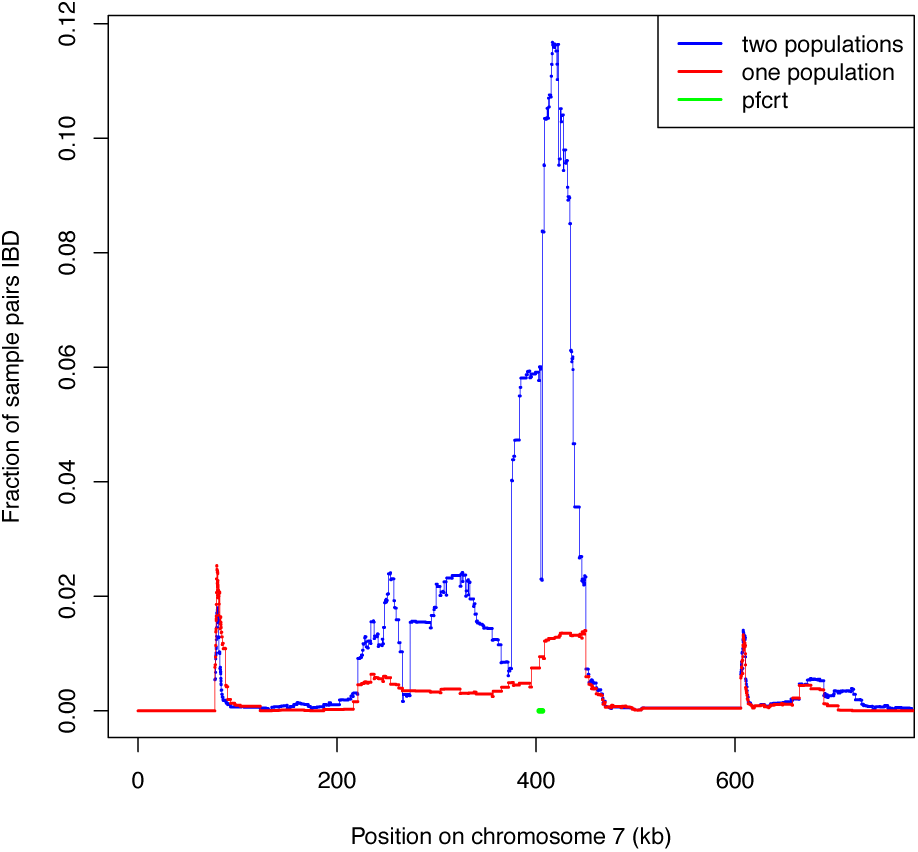
The fraction of sample pairs that are IBD along chromosome 7, where one sample is from Ghana and the other from Cambodia. Blue curve: IBD as reconstructed by hmmIBD correctly treating the samples as coming from two populations; red curve: IBD as reconstructed from a single, averaged population. (See S2 for details.)

Summary of average results based on IBD segments of artificially recombined field data with standard deviations in parentheses; full details can be found in Appendix S3.

## Acknowledgements

The authors wish to thank Jacob Garcia for providing estimates of complexity of infection for the Pf3K samples, as well as Caroline Buckee for advice and support, and Pierre Jacob and Patrick Rebeschini for helpful discussions.

